# Nanobody-antigen catch-bond reveals NK cell mechanosensitivity

**DOI:** 10.1101/386094

**Authors:** Cristina Gonzalez, Patrick Chames, Brigitte Kerfelec, Daniel Baty, Philippe Robert, Laurent Limozin

## Abstract

Antibodies are key tools in biomedical research and medicine. Their binding properties are classically measured in solution and characterized by an affinity. However, in physiological conditions, antibodies can bridge an immune effector cell and an antigen presenting cell, implying that mechanical forces apply to the bonds. For example, in antibody-dependent cell cytotoxicity, a major mode of action of therapeutic monoclonal antibodies, the Fab domains bind the antigens on the target cell, while the Fc domain binds to the activating receptor CD16 (also known as FcgRIII) of an immune effector cell, in a quasi bi-dimensional environment (2D). Therefore, there is a strong need to investigating antigen/antibody binding under force (2D), to better understand and predict antibody activity *in vivo.* We used two anti-CD16 nanobodies targeting two different epitopes and laminar flow chamber assay to measure the association and dissociation of single bonds formed between microsphere-bound CD16 antigens and surface-bound anti-CD16 nanobodies (or single domain antibodies), simulating 2D encounters. The two nanobodies exhibit similar 2D association kinetics, characterized by a strong dependence on the molecular encounter duration. However, their 2D dissociation kinetics strongly differ as a function of applied force: one exhibits a slip bond behaviour where off-rate increases with force; the other exhibits a catch bond behaviour with off-rate decreasing with force. This is the first time, to our knowledge, that catch bond behaviour was reported for antigen-antibody bond. We further exploit this property to show how Natural Killer cells spread differentially on surfaces coated with these molecules, revealing NK cells mechanosensitivity. Our results may also have strong implications for the design of efficient bispecific antibodies for therapeutic applications.

## INTRODUCTION

Antibodies are major research, diagnostic and therapeutic tools. These 150 kDa proteins can bind specifically most of natural and artificial targets (so called antigens). In mammals, after contact with a new antigen, highly specific and affine antibody proteins are produced by monoclonal B cells which are selected in germinal centers in a process called affinity maturation (1, 2). It was recently discovered that selection of high affinity antibodies occurs when B cells pull actively on their antigens, by exerting direct mechanical force on the antibody-antigen bond (3). Indeed, antigen-antibody bonds often act at cell-cell interfaces, for example between a pathogenic cell and an immune effector cell, including Natural Killer (NK) cells, during Antibody Dependent Cell Cytotoxicity (ADCC) or macrophages, during Antibody Dependent Cell Phagocytosis (ADCP), which leads to the destruction of the pathogenic cell by the immune cell (1). The functional contact established between NK cells or B cells and their target, the so-called immunological synapse, is highly organized by the actomyosin network and the physical forces it produces (4–7). The quality of the antibody binding is traditionally described by an affinity measured in conditions where one of the partner (antibody or antigen) is in solution; this parameter might not be completely relevant to describe their behaviour when tethered at surfaces and subject to mechanical disruptive forces, further referred to as “2D” environment (8).

The study of protein-protein interactions, like antigen-antibody, have been profundly renewed by the development of single molecule manipulation and measurements (9). These techniques measure interactions between complementary proteins tethered to opposite surfaces which are first put into contact and then separated. They have been successfully used to study: (i) unbinding force of biotin-streptavidin bond with Atomic Force Microscopy (10), (ii) anti Immunoglobulin-Anti-Ig kinetics with the Laminar Flow Chamber (11), (iii) biotin-streptavidin energy landscape of dissociation with the Biomembrane Force Probe (12). Bonds behave typically as slip bonds, whose lifetime decreases with applied force, as predicted by Bell’s law (13). However, catch bonds, whose lifetime increases with force, were initially discovered for physiological process such as bacterial adhesion (14) and selectins-mediated interaction between white blood cells and endothelial cells in response to infection (15). This behaviour has been later found in other systems including adhesion molecules such as cadherins and integrins and in the T cell receptor (16). However, to our knowledge, no catch bond has been described for antigen-antibody interaction (5).

The Laminar Flow Chamber (LFC) uses hundreds of microspheres conjugated to ligands and convected by a flow above complementary receptors immobilized onto a surface. At low flow velocity and low surface coated molecules density, it allows efficient ligand-receptor mechanical discrimination at the single bond level with the advantage of naturally multiplexed measurements (11, 17–19). Several original features of some antibody/antigen interactions were observed using LFC in this setting. For example, survival curves exhibited features of bond strengthening over the time after their formation (20); analysis of antibody/antigen association also revealed a non linear dependence of bond formation probability as a function of the duration of the molecular encounter between the reactive partners before bond formation, an observation questioning the definition of an association rate between surface tethered proteins (21–23). Whether these features are characteristic of many antigen-antibody bonds is important for a fundamental understanding of Ag-Ab interaction as well as for the technical validation of LFC measurements.

Nanobodies (aka single domain antibodies, sdAbs, or VHH) are antibody fragments derived from camelidae antibodies devoid of light chain. With a molecular weight of 15 kDa, and constituted of a single immunoglobulin domain, they can be used to target hidden epitopes or as elementary bricks to construct multispecific molecules (24). They can also circumvent limitations of conventional antibodies for certain diseases, by targeting cryptic conserved epitopes. Very recently, they were used as a library of cell-cell linkers for the engineering of multicellular aggregates (25). Due to their standardized monovalent format, a panel of nanobodies targeting the same antigen constitutes an ideal set to test the questions raised above. We have previously generated a set of nanobodies targeting the low affinity receptor CD16 (aka Fc*γ*Receptor III) expressed on NK cells and macrophages (26). Their on/off kinetics was measured in solution by Surface Plasmon Resonance (26). CD16, which binds the Fc fragment of conventional antibodies, is involved in ADCC and ADCP, so naturally subject to disruptive force generated within the immune synapse. Anti-CD16 nanobodies are surrogate Fc fragments which can form stronger bond than the Fc*γ*RIII-Fc fragment interaction, and that are dedicated to be coupled to another nanobody with a different specificity, in a bispecific construction (27). Such constructions, designed to be insensitive to CD16 polymorphism, were successfully tested to treat HER2 positive breast cancer with low HER2 expression resistant to the therapeutic monoclonal antibody trastuzumab (28). More generally, anti-CD16 nanobodies may serve as universal targeting moiety in various diseases (29) and their kinetic characterization under force would be a valuable information to select the most efficient binders in 2D settings.

In this work, we perform for the first time a comparative study of the association and dissociation kinetics of two nanobodies (named C21 and C28) targeting the same human antigen CD16 in the LFC. After insuring that conditions for single bond kinetics measurements were fulfilled, flow velocity was systematically varied. Association probability displays very similar behaviour for the two nanobodies, as a power law of the molecule interaction duration. The dissociation process shows a strengthening with time for the two nanobodies. However, the dependence of the initial off-rate with force strongly differs: one increases when force increases (slip bond), the other decreases (catch bond). This study identifies, for the first time to our knowledge, a catch bond behaviour for an antibody. We further show that NK cell spreading on nanobody-coated surfaces is more efficient when mediated by the catch bond nanobody as compared to the slip bond nanobody, implying that NK cells are are applying and sensing forces. Finally, NK cells adhesion under increasing shear force was markedly increased on surfaces coated with the catch bond nanobody. This work illustrates how the comparative use of antibodies which unbinding kinetics are well characterized under force can help deciphering complex cellular behaviours.

## MATERIALS AND METHODS

### Molecules and cells

Nanobodies C21 and C28 were previously generated after immunization of lamas with the recombinant human Fc*γ*RIIIB and selected by phage display as described in (26). GenBank accession number are: EF5612911 for C21; EF561292 for C28. Here C21 and C28, which both exhibit C-terminal c-Myc and 6 His tags were produced in *E. coli* and purified by TALON metal-affinity chromatography as previously described (26) (Fig. S7A). The transglutaminase-catalyzed biotinylation of the c-Myc tag was performed using the Biotin TGase Protein Labelling kit (Zedira, Darmstadt, Germany) following manufacturer instructions. After 1h incubation with biotinylation reagents at 22 °C, nanobodies were filtered using ZebaTM Spin Desalting Columns (ThermoFischer Scientific). Biotinylation of nanobodies was assessed by migration on gel using GelDoc ™ EZ Imager (Biorad, Hercules, California) for nanobodies bands visualization Western Blot using anti-His-HRP antibody (clone GG11-8F.3.5.1, Miltenyi Biotec, Paris, France) at 1/5000 and Streptavidin HRP at 1/2000 (ThermoFischer Scientific, Villebon-sur-Yvette, France) (Fig. S7B). Concentration of nanobodies were determined by measuring amine bonds in protein chains by infrared spectroscopy (Direct Detect Infrared Spectrometer).

Natural Killer NK92^hCD16^ cell line was used to perform cell adhesion experiments on nanobodies coated surfaces. NK92 cells were transfected to express a chimeric molecule containing the extracellular domain of human CD16 (Fc*γ*RIIIA-V158) and the transmembrane and intracellular domain of Fc*∊*RI*γ* as described by (30). Cells were cultured in RPMI 1640 + 10 *%* foetal bovine serum, (IL-2 Proleukin, Novartis, Bale, Switzerland) at 200 U/ml. Expression levels of CD16 were controlled once per week by flow cytometry using a fluorescent anti-CD16 (Phycoerythrin anti-CD16 human, clone 3G8, Biolegend, London, UK).

### Single bond kinetic measurements with the Laminar Flow Chamber

For laminar flow chamber (LFC) experiments with microspheres, glass slides were functionalized with biotin-conjugated anti-CD16 nanobodies as described before (18, 23). Briefly, slides were incubated successively with poly-L-lysine, glutaraldehyde, bovine serum albumine biotin, glycine, streptavidin (all products, Sigma Aldrich St Quentin Fallavier, France) and finally biotinylated anti-CD16 nanobodies at different concentrations. The detailed procedure is described in Supplementary Material. The nanobodies density on the surface at the various incubation concentrations was determined by fluorescence microscopy. For this purpose, surface functionalized with nanobodies were further incubated for 30 min with a fluorescently labelled anti-His-Phycoerythrin (anti-His-PE, clone GG11-8F.3.5.1, Miltenyi Biotec). The antibody is labelled in average with 1.5 PE group and binds the Histag of the nanobody. The detailed procedure for surface density measurement is described in Supplementary Material.

For microsphere functionalisation with recombinant CD16, 500 *μ*l of microspheres functionalized by toluenesulfonyl groups (Dynabeads M-450 Tosylactivated, ThermoFischer Scientific) of 4.5 *μ*m of diameter were rinsed in borate buffer 3 times. Then, 200 *μ*l of a solution of 0.5 *μ*g/ml anti Glutation-S-Transferase (anti GST) (Clone P1A12, Biolegend) were added to the microspheres resuspended in 300 *μ*l of borate buffer supplemented with BSA 0.1% and sodium azide 0.1% and the solution was incubated for 24 h at room temperature. Next, microspheres (40 *μ*l) were rinsed with PBS-BSA 0.2% and incubated with 10 *μ*l of a solution of 0.10 mg/ml of CD16 GST (human Fc*γ*IIIA GST tag recombinant protein (P01, Abnova, Taipei City, Taiwan) during 30 min with shaking. After this time, microspheres were cleaned with PBS-BSA 0.2% and directly used.

Single bond measurements were performed using a homemade automated Laminar Flow Chamber apparatus, composed of three mechanical systems coupled to an imaging system (23). Briefly, a glass slide coated with the nanobodies on the surface formed the bottom a multi chamber device with nine independent chambers used to test several densities of nanobodies on the same sample. The device was connected to one system that injects microspheres, another that controls the flow applied to the microspheres and the last one that regulates the temperature inside each chamber. Observation was performed using an inverted microscope equipped with a 20x/0.32 objective (1 pixel= 0.33 *μ*m) and images were recorded at a frame rate of 50 images/s using a camera (IDS). The temperature was set to 37°C.

Data were analysed as follows: the velocity of the microspheres was calculated on a time interval of 200 ms. The velocities of the sedimented microspheres (which correspond to the ones at molecular distance of the surface) were distributed around a peak *u_p_* ~ 0.54αG where a is the microsphere radius and G the shear rate (22). An interval of velocity was chosen around *u_p_* (Fig. S1B). The velocity should be within this interval in order to: (i) count the beginning of an arrest; (ii) count the travelled distance. On these velocity intervals, arrests of the microspheres were identified on the trajectories and counted (Fig. S1C). A microsphere was defined as arrested when its displacement *δx* was lower than 0.33 *μ*m during the defined time interval *δt* = 200 ms. The true arrest duration *d_ture_* was derived from the apparent arrest duration *d_app_* with the correction *d_ture_* = *d_app_* + *δt* – 2*δx*/*u_p_* (20). To analyse 2D association, the Binding Linear Density (BLD) was defined as the number of arrests divided by the travelled distance (23). In order to smoothen the data, the BLD were first fitted as a function of the velocity for a given density. Then, a series of velocities were chosen and the interpolated BLD values were used for further analysis (Fig. S1D). To analyse 2D dissociation, arrest durations were used to build the survival curves, i.e. the fraction of bonds still existing after time t.

### NK cells spreading experiments

For cell spreading experiments, uncoated *μ*-Slide 8 wells (Ibidi, Munich) composed of eight independent chambers were used. The surface coating with nanobodies was performed with 2 intermediate layers of BSA-biotin and streptavidin, before the deposition of monobiotinylated nanobodies (see Supplementary Material for details). Cell adhesion was monitored using Reflection Interference Contrast Microscopy (RICM), which is sensitive to cell-surface distance (31). Image acquisition starts immediatly after deposition of the cells in the devices. In order to determine the kinetics of spreading, several fields were selected and imaged cyclically during 10 min using a motorized stage (Physik Instruments). Elapsed time between two subsequent images on the same field was typically 20 to 30 s. After 10 min of cell incubation on the surfaces, about 20 to 30 fields were imaged both in transmission and reflection, in order to determine the proportion of adhering cells, their spreading area and the tightness of adhesion. Image analysis was performed to detect and measure adherent and non adherent cells on the coated nanobodies surfaces, and to distinguish them automatically from cell fragments. For this, images obtained sequentially in transmission and reflection, were exploited simultaneously using different home-made procedures. The detailed method is described in Supplemental Material. The kinetics of cell spreading was measured by segmenting cells on RICM sequences as described before (32). The area vs time curves were fitted with sigmoid function to extract a typical spreading time.

### NK cells laminar flow experiments

Two kinds of experiments with NK cells under laminar shear flow were performed. First, we measured the number and duration of adhesion events of NK cells freely moving in a shear flow on a CD16 nanobodies decorated surface. Uncoated μ-Slides IV0.4 (forming six independent channels) were coated with biotinylated anti-CD16 nanobodies as described for spreading experiments. 200*μ*l of a suspension of 800 000 cells per ml were injected in the device before each measurement. A second home-made model of automated laminar flow chamber device controlled a video camera and a syringe pump and applied successively shear stresses of 0.075 dyn/cm^2^, 0.3 dyn/cm^2^ and 0.6 dyn/cm^2^, while acquiring an independent video for each shear condition. Video analysis of cell trajectories was performed using the same algorithms than for microspheres described above and retrieved arrests lifetimes. Second, de-association of NK cells was also measured in different conditions. Using the same experimental set-up with a different automaton program, cells were injected in the chamber under a so-called “start flow” of 0.15 dyn/cm^2^ for 20 sec. Cells were then allowed to settle for 60 sec under a very low shear stress of 0.03 dyn/cm^2^(so-called “adhesion flow”), that still allowed to discriminate between adherent and no-adherent cells. Cells were then submitted to a series of higher shear stress, increasing by steps of 15 sec each as following: 0.2 dyn/cm^2^, 0.5 dyn/cm^2^, 1 dyn/cm^2^ and 2 dyn/cm^2^(so-called “de-adhesion flows”). For the de-adhesion analysis, number of adherent cells (N) was counted at the end of all the periods (N0, NI, NII, NIII, NIV and NV) (see Fig. 4). Proportion of adhering cells at each period (adhesion and de-adhesion) was determined by dividing the number of cells resting at the end of each period by N0 (or the total number of initially adherent cells).

## RESULTS

### Binding Linear Density and single bond assessment in Laminar Flow Chamber

To study the Binding Linear Density, each nanobody was incubated on the slides at 6-7 different concentrations ranging from from 0.004 to 0.125 *μ*g/ml, including a negative control without nanobody, leading to 6-7 molecular densities. For each coated surfaces, the shear rate in the LFC was set successively to 6 different values. The Binding Linear Density was plotted against nanobody surface density for each velocity condition, as shown in Fig. 1A (nanobody C21) and 1B (nanobody C28). For a given velocity, and in the range of selected densities, the BLD increases linearly with the molecular density, which indicates measure of single molecular bonds as multiple binding leads to saturation of the BLD. The data were fitted with an affine function, using a weight at each point corresponding to the error bar (most often linearity coefficient R>0.9). The interaction of the fitting line with the vertical axis represents the fitted non specific BLD. It is used to calculate the non-specific adhesion ratio r defined as the non-specific BLD divided by the BLD at a given condition.

**Figure 1:**
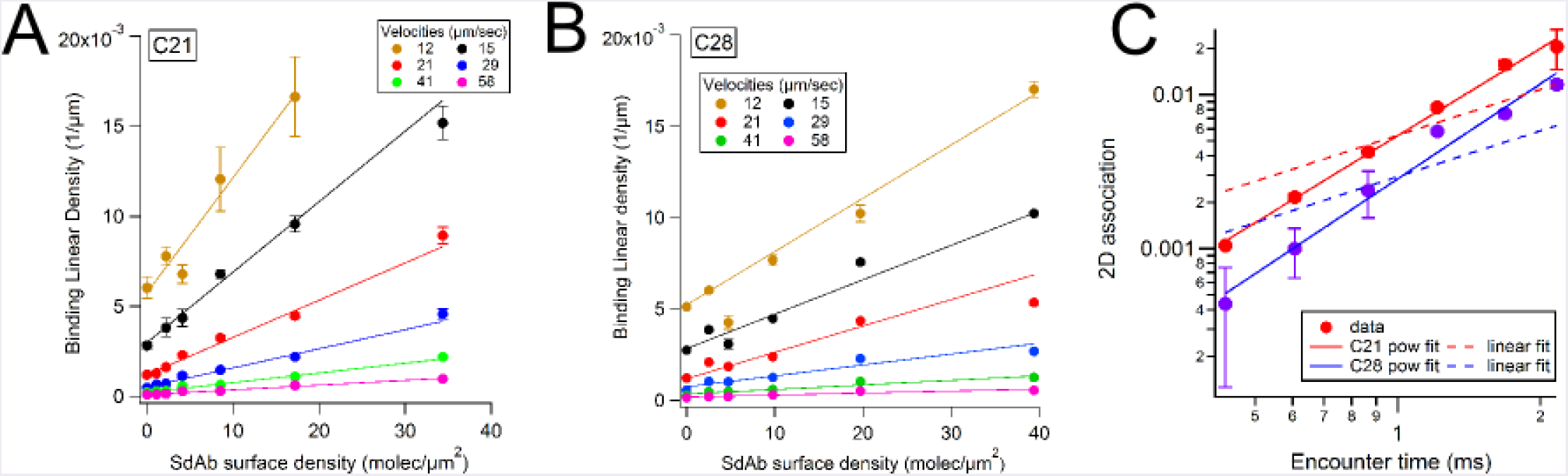
Binding linear density and 2D association measured with the laminar flow chamber. A, B). Binding Linear Density plots vs nanobody C21 (A) and nanobody C28 (B) surface density obtained at 6 velocity peaks *u_p_* of the sedimented microspheres. A linear fit of the data is presented for each *u_p_*. The error bars were defined as BLD divided by the square root of the number of arrests counted for the considered condition. C). Plot of the 2D association (corresponding to the slope of the BLD vs density linear fit, normalized by the molecular length L=25 nm (see Fig. S6) as a function of the encounter time (=*u_p_*/L) for C21 (red) and C28 (blue). The error bars were calculated by the variation of the slope when considering the linear fit of BLD vs density line, obtained on a narrower density range (by removing the highest density). Data were fitted to a power law (plain line) or a linear law (dashed line).

At a given experimental condition, the survival curve for specific arrests was built by subtracting from the total survival curve a fraction r of arrests following the non-specific survival distribution, i.e. measured in the absence of nanobody (20). The corrected survival was calculated as 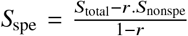. On fig. S2, the resulting curves are presented for 5 different velocity intervals and 3 different incubation concentration of nanobody, corresponding to 3 molecular densities. Each curve represents at least 150 arrests and are restricted to ratio *r* > 0.65. For given nanobody and density, the curves superimpose, demonstrating that the dissociation kinetics is independent of the density in this range, ruling out multiple binding which leads to lower dissociation. Taken together with the linear dependence of BLD on density, this is a strong assessment for single bond measurements (17, 23).

### Molecular Association

The 2D association was defined for each velocity as the slope of the BLD vs density line divided by the molecular length L. The normalization by L accounts dimensionally for the effect of molecular length in estimating the number of molecular encounters. A more precise modeling involves complete brownian dynamics simulations and the possible rotation of the molecules (22, 23). On Fig. 1C, the 2D association A_2D_ are represented as a function of the molecular encounter time *t*_enc_, defined as the ratio of molecular length L and velocity *u_p_*. The 2D association is well represented by a power law : 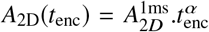 with *t*_enc_ in ms. Values of the fitting parameters are reported in Table 1. A tentative linear fit (shown as dashed line in Fig. 1C) emphasizes the finding that the association does not scale linearly with the encounter time. This was already observed in LFC for conventional antibodies (21–23).

**Table 1:**
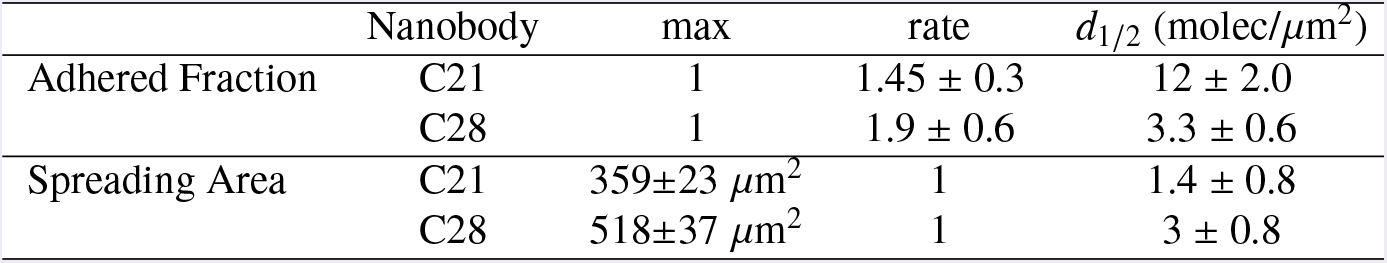
Summary of time and force dependent kinetics parameters of anti-CD16 nanobodies measured by laminar flow chamber.

### Molecular Dissociation

The survival curves displayed in Fig. and S2 exhibit a non linear shape in semi-log representation, indicating the involvement of different time scales in the dissociation process (17, 20). Curves of Fig. 2 A, B were fitted between 0 and 5 s, using the empirical equation:

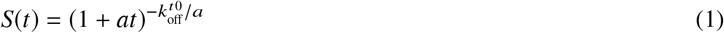

where 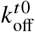 is the initial dissociation rate (in s^−1^) and *a* the rate of bond strengthening (in s^−1^), as applied earlier for conventional antibodies (20). Curves of Fig. 2 A, B also evidence the dependence of the survival on the external force applied to the bond through the flow. The force was proportional to the velocity as *F* (pN) = 1.25 *u* (*μ*m/s) (17, 20). Therefore, the parameters 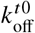 and *a* are force dependent. The average values of 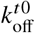 and *a*, calculated from the fits of survival curves obtained at 3 molecular densities, are displayed on fig. 2 C,D. The error bars are the standard deviation calculated with the 3 densities. Nanobody C21 exhibited a clear increase of the initial off-rate when force increases, which is characteristic of a slip bond. On the contrary, for C28, initial off rate decreased when force increases, which is characteristic of a catch bond. The strengthening parameter *a* was roughly independent of force for C21 and decreased with force for C28. 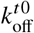 was fitted with Bell’s equation (13) : 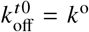.exp(*F*/*F_k_*). *k^O^* represents the off-rate at zero force; *F_k_* represents the typical force above which the off-rate becomes force dependent. The strengthening parameter *a* was simply fitted with an affine law *a* = *a^O^*.(1 + *F*/*F_a_*). While this dependence could be justified with some arguments of friction on the energy landscape of the interaction (P. Bongrand, personal communication), we use it here simply as a functional dependence in order to calculate the off-rate at any force and time. Values of the fitting parameters for both 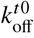 and *a* are reported in Table 1. These parameters allow to calculate the dissociation rate for any applied force and maturation time, using Eq. 1 (Fig. S3). Interestingly, the ratio of the off-rates shows that for durations above 1 s or applied force above 20 pN, C28 was more stable than C21 (Fig. 2E).

**Figure 2:**
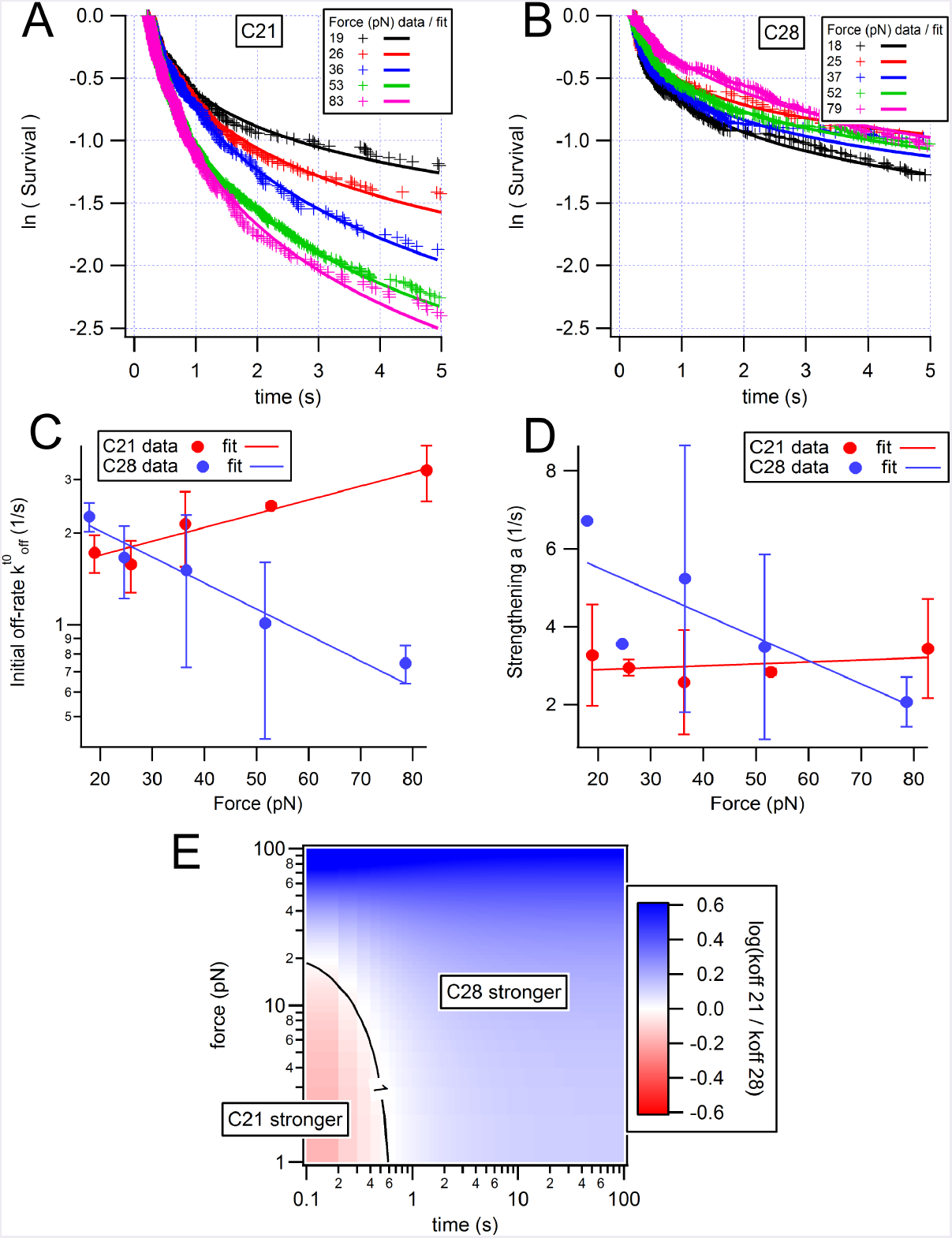
Analysis of dissociation of nanobodies C21 and C28 from CD16. A-B) Survival curves for 125 ng/ml nanobody incubation concentration at various applied forces (in pN). Each curve was fitted with equation 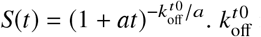 is the initial dissociation rate and *a* the rate of bond strengthening. C-D) These rates are represented as a function of the force and fitted with Bell’s law 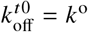.exp(*F*/*F_k_*) or an affine law *a* = *a^o^*.(1 + *F*/*F_a_*). The error bars correspond to the standard deviation of the fit parameters obtained for three different incubation concentrations (31, 62, 125 ng/ml) of nanobody. E) Ratio of calculated off-rates as a function of applied force and bond lifetime.

### Cellular spreading measured by RICM

To assess the effect of the two different molecular kinetics at cellular scale, the spreading of NK92 cells expressing CD16 on surfaces coated with either C21 or C28 was studied using RICM. The surface density of nanobodies was systematically varied between 1 and 100 molec/*μ*m^2^, as measured after each experiment, using the procedure described in Fig. S4A,B. The state of NK cells in terms of CD16 expression was controlled regularly by flow cytometry Fig. (S4C). Their spreading capacity was assessed regularly by measuring their spreading area and reflectivity on control surfaces coated with a conventional anti-CD16 antibody (S4D, E). The fraction of spread NK cells was measured after 10 min of engagement on the surface, by counting the number of cells displaying an adhesion patch by RICM divided by the number of cells visible by transmission, as described in details in Supplementary Material. The spread fraction increases with antibody surface density, with the fraction being larger for C28 at most densities (Fig.3A). The spread fraction as function of the nanobody molecular density *d* was fitted with a Hill equation 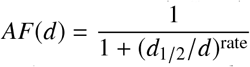. The fitted parameters *d*_1_/_2_ and rate are reported in Table 2. The value of half density *d*_1_/_2_ determined for nanobody C28, *d*_1_/_2_ = 3.3 ± 0.6, was 4-fold lower than that determined for nanobody C21, indicating that NK cells adhere on lower densities of C28 than C21. The spreading area of cells after 10 min of engagement was also measured as a function of nanobody coverage (Fig.3B). A fit with Hill equation was applied by fixing the rate to 1 and fitting the maximal area yielding 359±23 *μ*m^2^ and 518±37 *μ*m^2^ for C21 and C28 respectively. Finally, the reflectivity of RICM images was also used to assess the distance between the basal membrane of NK cells and the nanobody-coated surface. Indeed, low grey level can be used as a proxy for short membrane-surface distance (31). This distance decreased with antibody surface density, and was smaller for C21 at most of the densities (Fig.3C). The kinetics of spreading was also recorded (Fig. S5). There was no significant difference between the duration of spreading on C21 and C28, tested at various surface densities.

**Figure 3:**
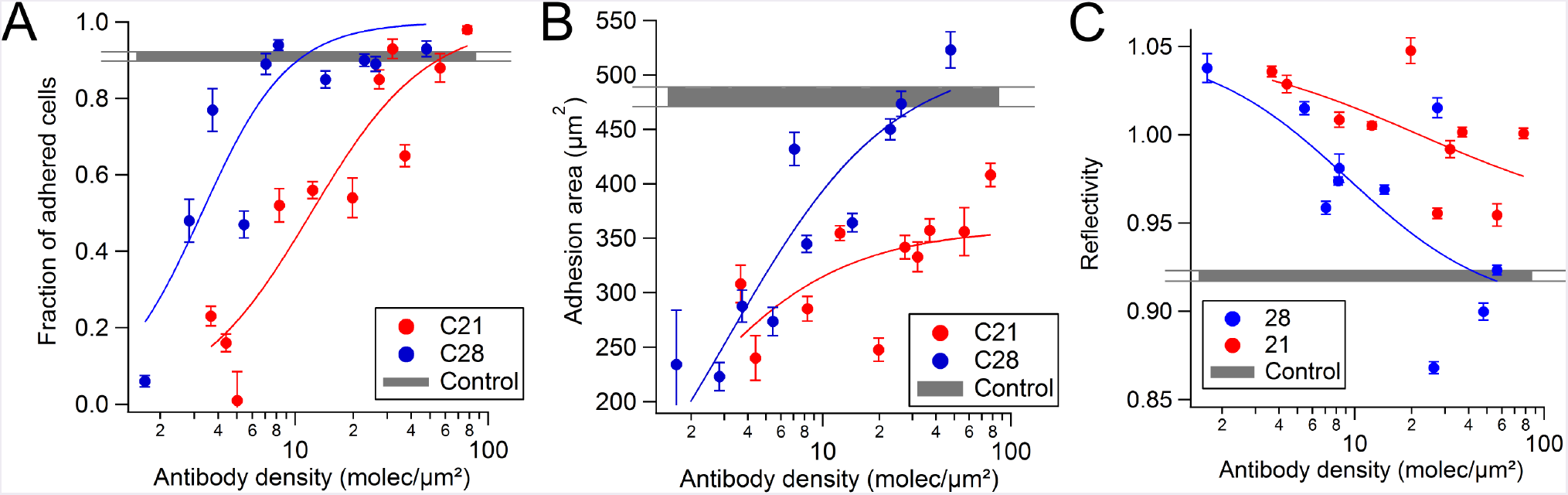
Adhesion of NK cells on nanobodies coated surface measured by RICM. A) Plot of the fraction of spread cells in function of nanobody density. B) Plot of the spread area as a function of nanobody density. C) Reflectivity signal of adhered cells, providing an estimate of the tightness of adhesion, as a function of nanobody density. In all experiments, controls correspond to cells spread on surfaces coated with conventional anti-CD16 antibody (see Fig. S4). Each point represents the pool of 4 separate experiments with at least 100 cells. Error bars are SEM.

**Table 2:**
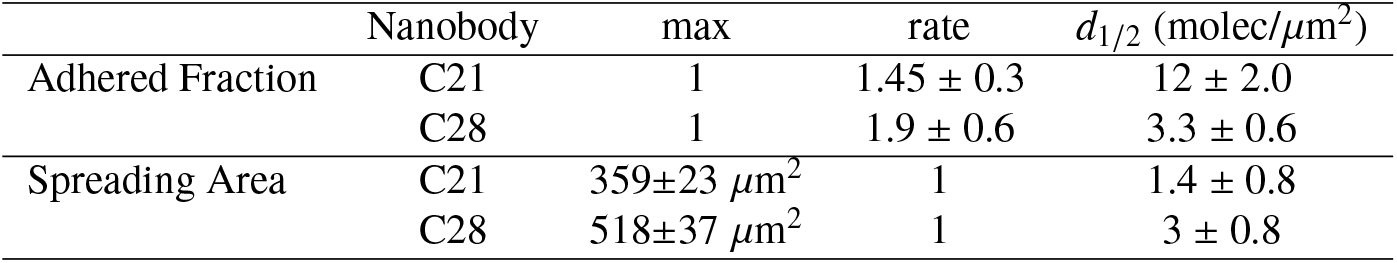
Summary of adhesion parameters of NK cells on anti-CD16 surfaces measured by RICM. A Hill equation 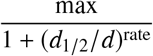 is fitted to the data to describe their dependence on nanobody surface density.

### Cellular transient adhesion and de-adhesion

To quantify further the adhesion of NK cells on nanobodies coated surfaces, we measured cell adhesion in the Laminar Flow Chamber. As C21 and C28 survival curves superimposed in all shear rate tested, transient adhesion of NK-92 cells on anti-CD16 coated surfaces does not show any difference between the adhesive capacity of C21 and C28 (Fig. S6). These results show that the difference in off-rate kinetics measured at the molecular scale is not visible at the cellular scale in transient adhesion experiments. It may be hidden by the formation of multiple bonds during the process.

To assess whether the off-rate kinetics plays a role for cells at a longer time scale, in line with the above observations concerning spreading, we let the cells adhere in the flow chamber several seconds before applying a series of flows of increasing shear rates (Fig. 4). Clearly, cells adhering on C28 resist better to the detachment force than cells adhering on C21, indicating that a duration of several seconds of engagement is required to observe the catch-bond effect of C28.

**Figure 4:**
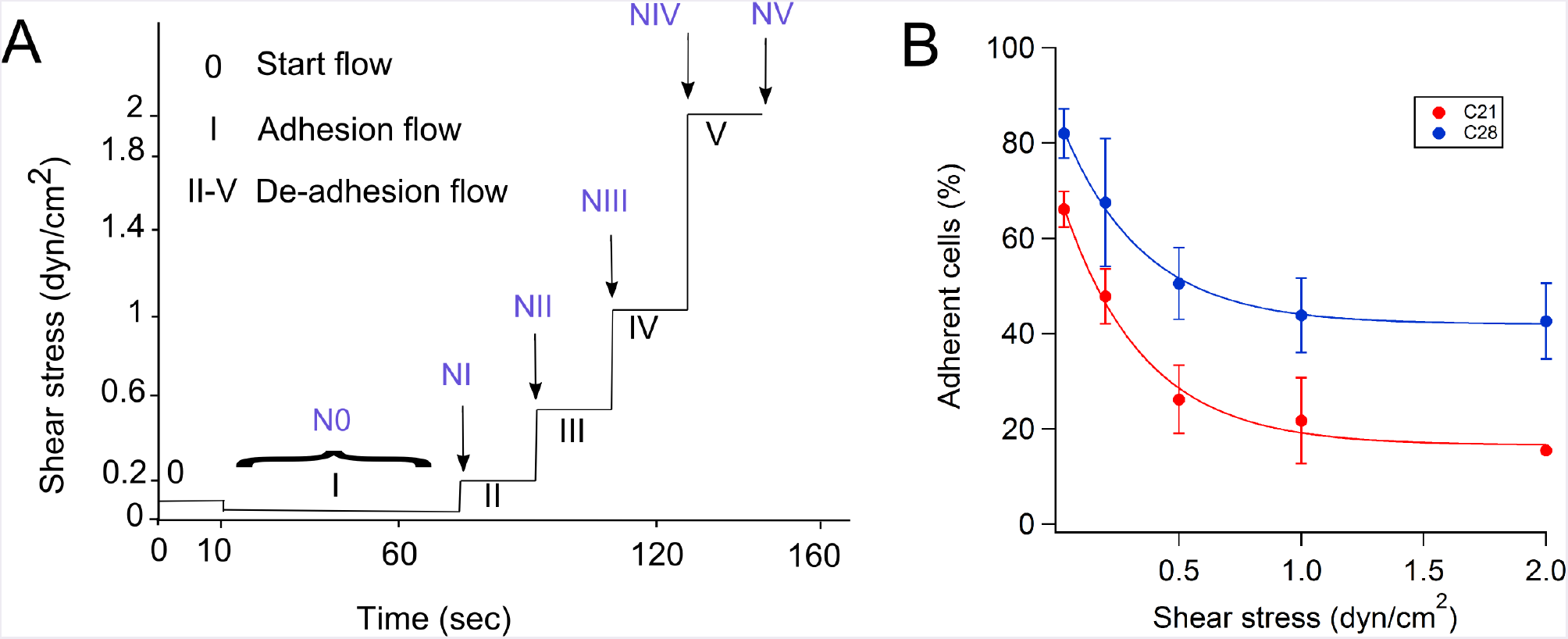
Detachment of cells adhering on nanobodies coated surfaces by a flow. A) Time sequence of the shear stress imposed on adhering NK cells in the laminar flow chamber. Most of the cells adhere to the surface during the period I (adhesion flow), and de-adhere during force steps II-V. B) Fraction of attached cells on nanobody coated surfaces at the various imposed shear stress. Nanobody coated densities were between 6.5 and 15 molecules/*μ*m^2^. Points represents mean values of two-three independent experiments. Number of total adherent cells detected were > to 50 for both nanobodies. Error bars are the standard deviations.

## DISCUSSION

The purpose of this study was to dissect the association/dissociation mechanisms between antibody fragments such as nanobodies and their antigen in order to identify new criteria in the perspective of designing nanobodies-based therapeutics. By measuring and comparing the binding of two nanobodies on the same antigen, we have evidenced comparable association and different dependence on the force of the dissociation. The Laminar Flow Chamber is a method of choice for rapid measurement of both association and dissociation kinetics of ligand-receptor bonds tethered at surfaces. The criteria of single bond assessment is very stringent, whereas alternative single bond techniques like AFM often rely only on a maximum of 10% of binding events observed (33). Applied flow limits the encounter duration between receptor on the microsphere and ligand on the underlying surface to the millisecond range. As a consequence, the external part of the energy landscape is probed, as it was shown for the biotin-streptavidin bond (9, 17). Therefore, the results reported here concerning the initial off-rate may not be valid for deeper internal parts of the energy landscape. Conversely, the technique allows to precisely control the time of bond formation in the millisecond range. This has two advantages: first, the interaction duration between the reactive partners can be varied and the resulting bond formation measured (23); thus, we were able to show that, as already observed for conventional antibodies, the 2D association varies non-linearly with the interaction duration (21–23). Second, bond maturation could be observed and quantified through the strengthening rate *a* (20). Nanobody-antigen bonds actually reinforced with time on the second timescale, as previously observed for conventional antibody-antigen bond (20). Interestingly, other immune interactions probed with LFC, like T Cell Receptor - peptide Major Histocompatibility Complex (TCR-pMHC), exhibit rather slower strengthening (P. Robert, unpublished data), suggesting that these observations are not an artefact due to the method. Nevertheless, further efforts should be undertaken to support the concept of bond maturation, through new development in the LFC, like variable flow, currently under test. Overall, our results emphasize that despite their small size, nanobodies exhibit complex association kinetics with their antigen, consistent with measurements on conventional antibodies.

The aforementioned technical limitations do not affect the comparative study presented here for several reasons. First, the dependence on encounter time of the 2D association is very similar for the two nanobodies, with exponent differing of less than 10%. This rules out the possibility of an artefactual difference in dissociation caused by significant difference in association. Additionally, it was described that the epitopes recognised by the two nanobodies are different, but closely located since both epitopes are shared with mAb 7.5.4, (26). As 2D association depends on the distance between molecules, similar on-rate favours the hypothesis of closely located epitopes with comparable molecular chain length L for the chains obtained with the two nanobodies in our setting (23).

The on/off kinetics of C21 and C28 have been measured previously using surface plasmon resonance (SPR) with diffusing nanobodies binding CD16 tethered to surfaces (26). Off-rate in solution (3D off-rate) was found to be 2.8 × 10^−3^ s^−1^ for C21 and 3.4 × 10^−3^ s^−1^ for C28. In this study, we find values of k^o^, the initial off-rate at zero force, about 1000 larger for both C28 and C21. THis discrepancy was already observed in the LFC for kinetics of antibodies or TCR-pMHC (11, 19). We attribute this discrepancy to the short encounter duration imposed by the flow, leading to the measurement of dissociation in an early state of the bond (23). This is however consistent with the bond strengthening. For example, after 100 s, we predict an off-rate at zero force of 5 × 10^−3^ s^−1^ for C21 and 4 × 10^−3^ s^−1^ for C28 (Fig. S3 A, B). Previous AFM studies showed a satisfying correlation between the 2D off-rate extrapolated at zero force (*k^o^*) and 3D off-rate as measured with SPR (34, 35). However, our results show that Bell Force are strongly different: *F_k_* ~100 pN for C21 corresponds to a potential width of 0.04 nm in the energy landscape, likely related to a stiff bond (36). For C28, *F_k_* -51 pN, which clearly shows a catch bond behaviour, as based solely on the survival curves. One should however consider also the strong reduction of BLD for high velocities (force), which may be the consequence of a selection in measured bonds. Concerning the association, the values of k_on_ provided by SPR measurements were 2.9 × 10^5^ M^−1^.s^−1^ for C21 and 0.4 × 10^5^ M^−1^.s^−1^ for C28. The conversion of our 2D association into a 3D k_on_ requires several assumptions on molecular length and flexibility (23). Qualitatively, C21 associates faster than C28 in 2D or 3D.

Our finding are particularly interesting in the perspective of designing bispecific antibodies used in therapeutics (27). For generating single domain antibody based bispecific antibodies (bsAbs), the binding properties of those anti-CD16 might be of outmost importance but the basis for choosing the best binder remains elusive. We have previously generated two anti-CEA bsAbs using a common anti-carcinoembryonic antigen (CEA) nanobody and either C21 or C28 (37). Interestingly, while the C21-based bsAb bound more efficiently to CD16 expressing cells by flow cytometry, probably reflecting the difference of dissociation constant K_*D*_, their ability to activate NK cells were very similar as evidenced by IL2 secretion assays and *in vitro* ADCC assays. Thus, while the accessibility of the CD16 epitope when displayed on the cell surface might clearly be a relevant consideration, these results suggest that a choice solely based on apparent affinity might be restrictive. C21-based bsAb was the chosen candidate for further resource and time-consuming animal studies (28, 37). However, our 2D measurements indicate here that C28 should exhibit a stronger resistance to force than C21. This is likely to be the case in the NK immune synapse, therefore indicating that C28 may be a better choice. Whether this parameter has an influence in the particular environment of the immune synapse deserved to be further investigated.

In the recent years, mechanical forces have been shown to play a central role in the immune system, for example with mechanotransduction, during cell migration or immune cell-cell interaction (38). This was specially studied in the case of the recognition of the T cell receptor with the pMHC, which was proposed to function as a catch-bond (39, 40). Much less is known about the mechanical response of antibodies and their possible physiological role. The T-cell and NK cell synapses exhibit strong ressemblance including the role of integrins (41), actin organisation and depletion for cytotoxic vesicle release (42), actin retrograde flow (6). Based on literature and our own experience with T-cells (32, 43), we hypothesize that the NK cell synapse is also exercing and sensing force. Our cellular experiments show that NK cells engage an immune synapse on anti-CD16 coated surfaces, for sufficiently high densities of antibodies. This does not require additional integrin ligands. It is likely that this process involves the cell pulling on the bond, and that C28 offers a better resistance than C21. Using the calculated ratio of the off-rates (Fig. 2E), we speculate that the force maybe above 10 pN and the duration of the pulling beyond 1 s. This is also consistent with the observation that C28 provides a larger maximal spreading area (Table 2). While much experimental and theoretical work will be required to establish a more quantitative link between the molecular and cellular scale, as attempted recently in the case of the TCR (44), or selectins in biomimetic systems (45), we show here the strong potential to use carefully force-characterized nanobodies as probes for deciphering cell mechanical behaviour.

## AUTHOR CONTRIBUTIONS

CG carried out all experiments and most of the analysis. PR and LL designed the research, supervised the experiments and the analysis. PC, BK and DB contributed nanobodies and cells. CG and LL wrote the article.

## ACKNOWLEDGMENTS

We thank M. Biarnes-Pelicot for help with cell culture and flow cytometry, D. Touchard for complementary de-adhesion experiments and P. Bongrand for critical reading of the manuscript.

## SUPPLEMENTARY MATERIAL

### Nanobody-antigen catch-bond reveals NK cell mechanosensitivity

Cristina Gonzalez, Patrick Chames, Brigitte Kerfelec, Daniel Baty, Philippe Robert, Laurent Limozin

### Supplementary Methods

#### Surfaces preparation for molecular measurements with the laminar flow chamber

For laminar flow chamber (LFC) experiments, glass slides of 75x25 mm^2^ (VWR) were rinsed twice with ethanol 98% and distilled water, then deposited 10 min in a solution containing 2/3 H_2_SO_4_ at 93-98 % and 1/3 of H_2_O_2_ at 50 % (both Sigma-Aldrich) then rinsed thoroughly with deionized water. Negative charged glass slides were incubated 10 min with a solution of 100 *μ*g/ml of polylysine (Poly-L-lysine hydrobromide 150000-300000 kDa, Sigma-Aldrich) in phosphate buffer 0.01 M pH= 7.4 with 0.01% azide. Slides were subsequently washed with PBS and incubated 10 min with 25 mg/ml of glutaraldehyde in borate buffer (H_3_BO_3_ + H_2_O) 0.1M pH=9 with 0.01 % azide. Amine groups of polylysine make covalent bonds with one of the aldehyde groups of glutaraldehyde. After washing with PBS, another incubation of 10 min with 100 *μ*g/ml of BSA biotin (Sigma-Aldrich) in PBS was performed. Glass slides were washed with PBS and incubated for 10 min with a solution of 0.2 M glycine in PBS + 0.1% BSA for neutralization of remaining free aldehyde groups. After washing with PBS, slides were incubated for 30 min with 10 *μ*g/ml of a solution of streptavidin (Sigma-Aldrich) in PBS. Finally, after washing with PBS, slides were deposited on the bottom of the LFC and 100 *μ*l of biotinylated nanobodies were incubated for 30 min at various concentration in each compartment, before a final rinsing with PBS.

#### Measurement of surface density of antibodies

Samples were imaged using a microscope Observer (Carl Zeiss) equiped with an objective 20x/0.8, a 200 W light source (Lumen200, Prior) set at 10% power and an additional neutral filter (transmission 30%) to reduce photobleaching. Illumination aperture was set to 0.95. Fluorescence was excited and collected with the following filterset: EX 546/12 nm - BS 560 nm - EM 575-640 nm. Images were recorded, using an Andor iXon camera and Micro Manager software, at different exposure times (50, 100, 200, 500 ms) depending on the fluorescence intensity of the sample, in order to optimize the signal. 10-20 fields were imaged for each sample. For the analysis, a region of interest (ROI) was defined for all images using Image J giving mean intensity values and the standard deviation for all the ROI of each image (Fig. S7C). From this mean value, the intensity given by the offset of the camera was removed and the result was divided by the exposure time. To retrieve the surface density of fluorescent molecules from the intensity, a calibration was performed by measuring the fluorescence of a known amount of fluorescent antibody in a 10 *μ*m thin volume (22) (Fig. S7D). The relation between surface density of antibody and incubation concentration was finally determined (Fig. S7E).

#### Surface and cell preparation for spreading experiments

Uncoated *μ*-Slide 8 wells were functionalized with single domain antibodies as follows: 100 *μ*g/ml BSA biotin (Sigma Aldrich) was deposited directly on the device and incubated 30 min. Then, devices were rinsed with PBS and incubated 30 min with 10 *μ*g/ml streptavidin (Sigma Aldrich) in PBS. Biotinylated nanobodies C21 or C28 were incubated at various concentrations during 30 min and devices were finally rinsed with PBS before cell deposition. A positive control was performed by replacing the nanobody by a conventional anti-CD16 biotinylated mAb (clone 3G8, Biolegend). nanobodies density on surface was measured by fluorescence as described above. Before each experiment, 20.000 cells were collected from culture flasks, centrifuged 5 min at 1500 rpm, re-suspended in 200 *μ*l of PBS-BSA 0.2 % and kept 10 min in Eppendorf tubes at 37 °C, before deposition in the device which was previously heated at 37 °C.

#### Image analysis procedure to determine adherent and non-adherent cells

Using Fiji distribution of ImageJ, Reflection (RICM) images were normalized by the background (to obtain reflectivity) and segmented as described previously (32). Briefly, a variance filter with a radius of 8 pixels or 1.6 *μ*m was applied to the reflectivity image, followed by a threshold at comprised between 0.002 and 0.008. The Analyse Particle plugin of ImageJ was then applied to define Regions of Interest (ROI) with a minimal area (fixed to 1000 pixel or 40*μ*m^2^, in order to remove small defects on images) and a minimal circularity fixed to 0.1. Two examples of normalized RICM images and ROI are shown on Fig. S8B and D. The same procedure was applied to segment cells from transmission images; the radius of variance was fixed to 5 pixel (or 1*μ*m), the minimal area fixed to 2000 pixel or 80*μ*m^2^ (higher than RICM images as in this case we focus on cells selection, not in adhesion area) and the minimal circularity fixed to 0.3. Transmission images with the ROI are shown on Fig. S8A and C.

Coordinates, area, mean and standard deviation of the intensity of all the ROI in reflection and transmission were measured using Fiji and transfered to Igor Pro software (Wavemetrics). A second threshold of size was made in order to remove cells fragments. ROIs with an area below 3500 pixels (140 *μ*m^2^) in transmission were removed except if the adhesion area (RICM) was above 3000 pixels (120*μ*m^2^). Based on the reflectivity properties on the negative control (no adherent cells) and on the positive control (almost all adherent cells), ROI were divided into 4 populations (P1, P2, P3, P4) in order to distinguish adherents and non adherent cells as shown in Fig. S9. P1 are ROI detected in reflection but not in transmission, corresponding to very adherents cells. P2 are ROI detected both in reflection and transmission with mean reflectivity below 1.07 and sd reflectivity below 0.06, corresponding to adherent cells. P3 are ROI detected in transmission but not in transmission, with mean reflectivity values between 1.02 and 1.07 and sd reflectivity values below 0.06, corresponding to non adherent cells. ROI corresponding to non adherent cells show as white patches in reflection and detected as ROI. To account for that, the population P4 was defined as ROI which appeared in transmission and reflection with the same reflectivity values as P3 (mean reflectivity values between 1.02 and 1.07 and standard deviation reflectivity values below 0.06) corresponding to non adherent cells. Once cells were classified into the 4 populations, adhesion cell fraction was calculated as 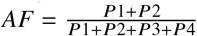.

Mean adhesion area and SEM from cells P1 and P2 (adherent cells) were calculated. To quantify the tightness of adhesion, mean and SEM reflectivity from cells P1 and P2 were calculated. For kinetics adhesion experiment, ROI were detected from reflection and their adhesion area was measured. In Igor Pro, knowing the position of the cells on the images, a criteria of minimal distance between the cells of different images was established and allowing individual cells to be tracked over all the stack of RICM images (32). Elapsed time between images was saved in metadata folder and used to track the adhesion area of cell over the time.

**Figure S1:**
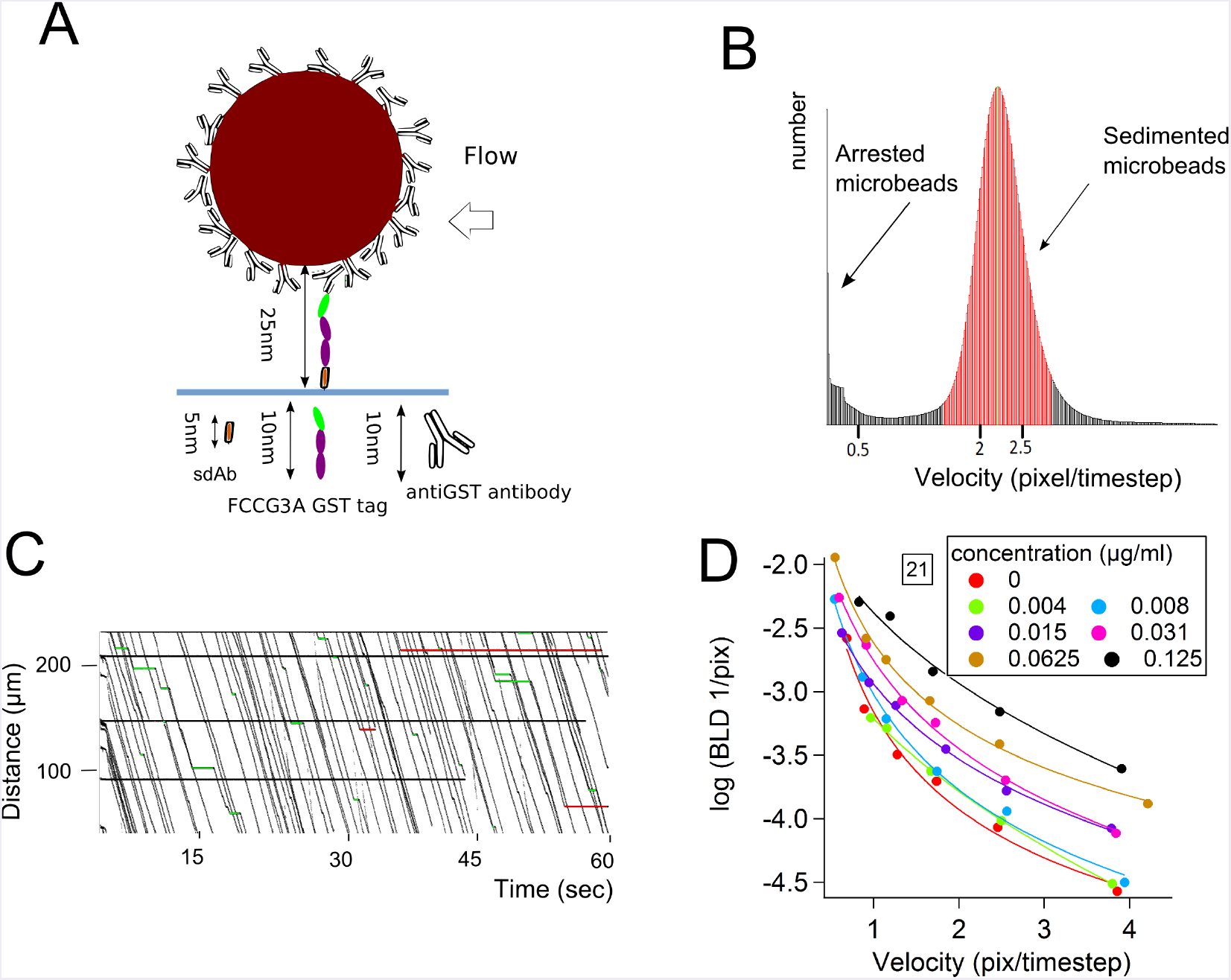
Laminar Flow Chamber for single bond kinetic measurements. A) Schematic representation of the strategy used to measure nanobodies-antigen interaction. Fc*γ*RIIIA (CD16) is coated to the microsphere via the anti-GST antibody and nanobodies are on the functionalised surface. Approximate length of all molecules is represented. Due to random orientation of the anti-GST antibody bound to the microsphere surface, an average length of 10 nm is considered. Total length of the molecular chain L=25 nm is represented and used to calculate, at a given shear rate, the molecular encounter time before bond formation and the force applied before bond rupture. B) Velocity histogram showing the peak of the arrested microspheres and the peak of the sedimented microspheres. C) Typical set of microsphere trajectories. Arrested microspheres are represented with a straight bar in green when the duration of the arrest is known and in red when is unknown. D) Interpolation of measured BLD as a function of microsphere velocity for each incubation concentration. Each data point corresponds typically to 4 independent experiments.

**Figure S2:**
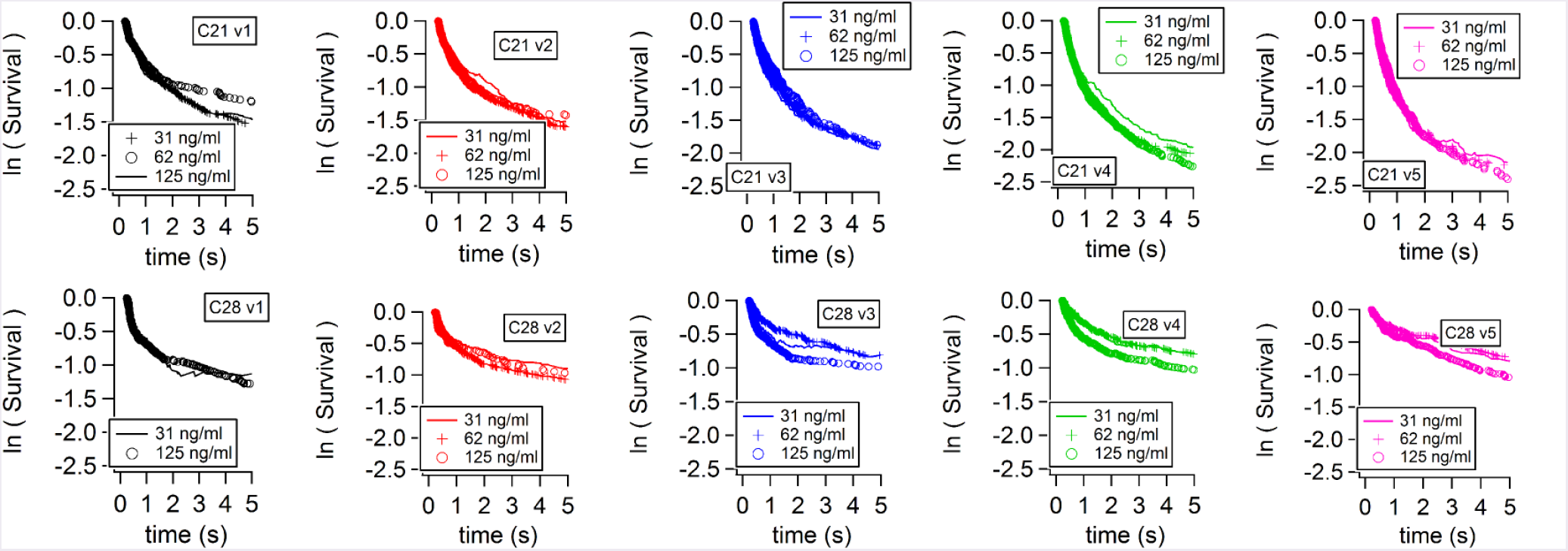
Superimposed survival curves for single bond assessment. Specific survival curves for nanobodies C21 (top row) and C28 (bottom row). Nanobodies were incubated at concentrations at 31, 62, 125 ng/ml and microsphere velocities were measured at 15, 21, 29, 41, 58 *μ*m/s. Curves superimposition at various molecular density of nanobody show that the dissociation kinetics do not depend on density, strongly supporting the measurement of single antibody-antigen bonds.

**Figure S3:**
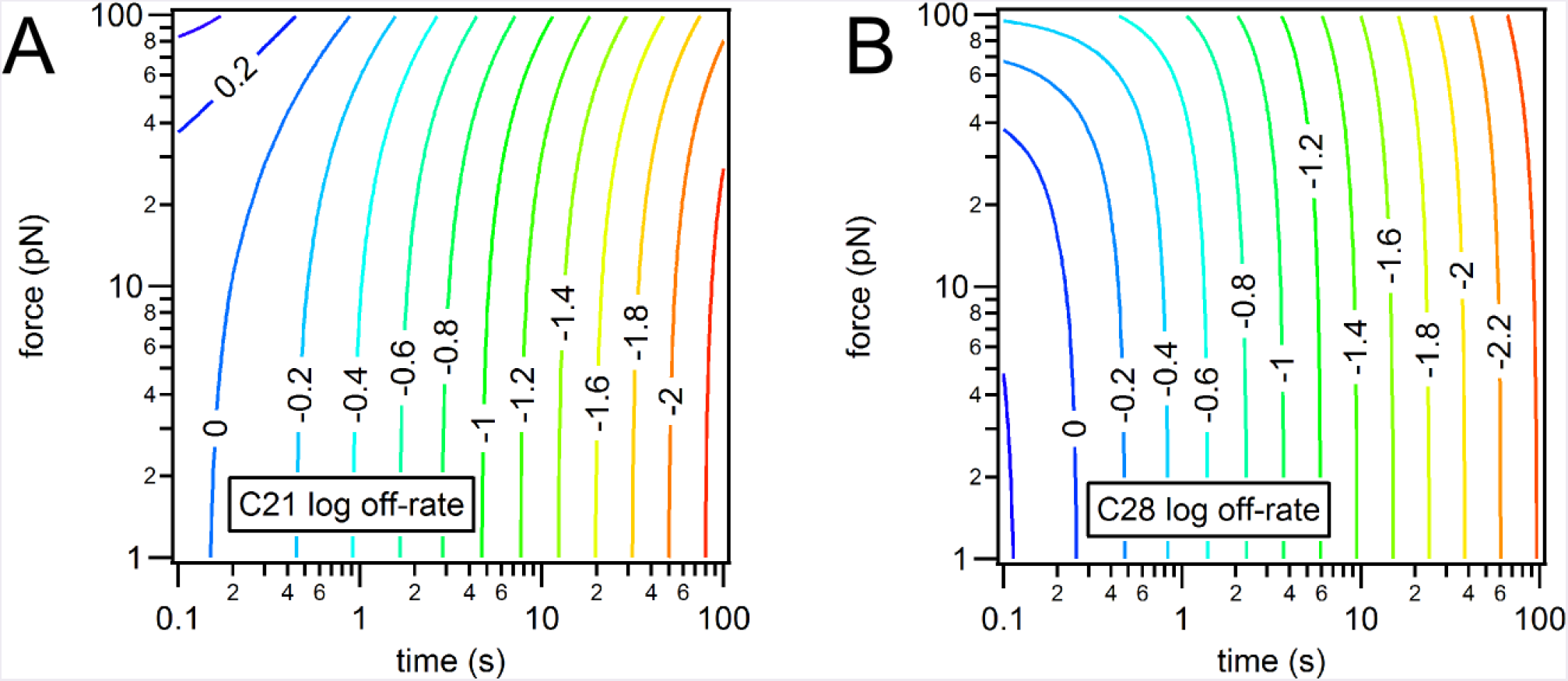
Logarithm of off-rates for C21 (A) and C28 (B) as function of applied force and lifetime of the bond. Values are calculated using measured parameters from Table 1 and Eq. 1.

**Figure S4:**
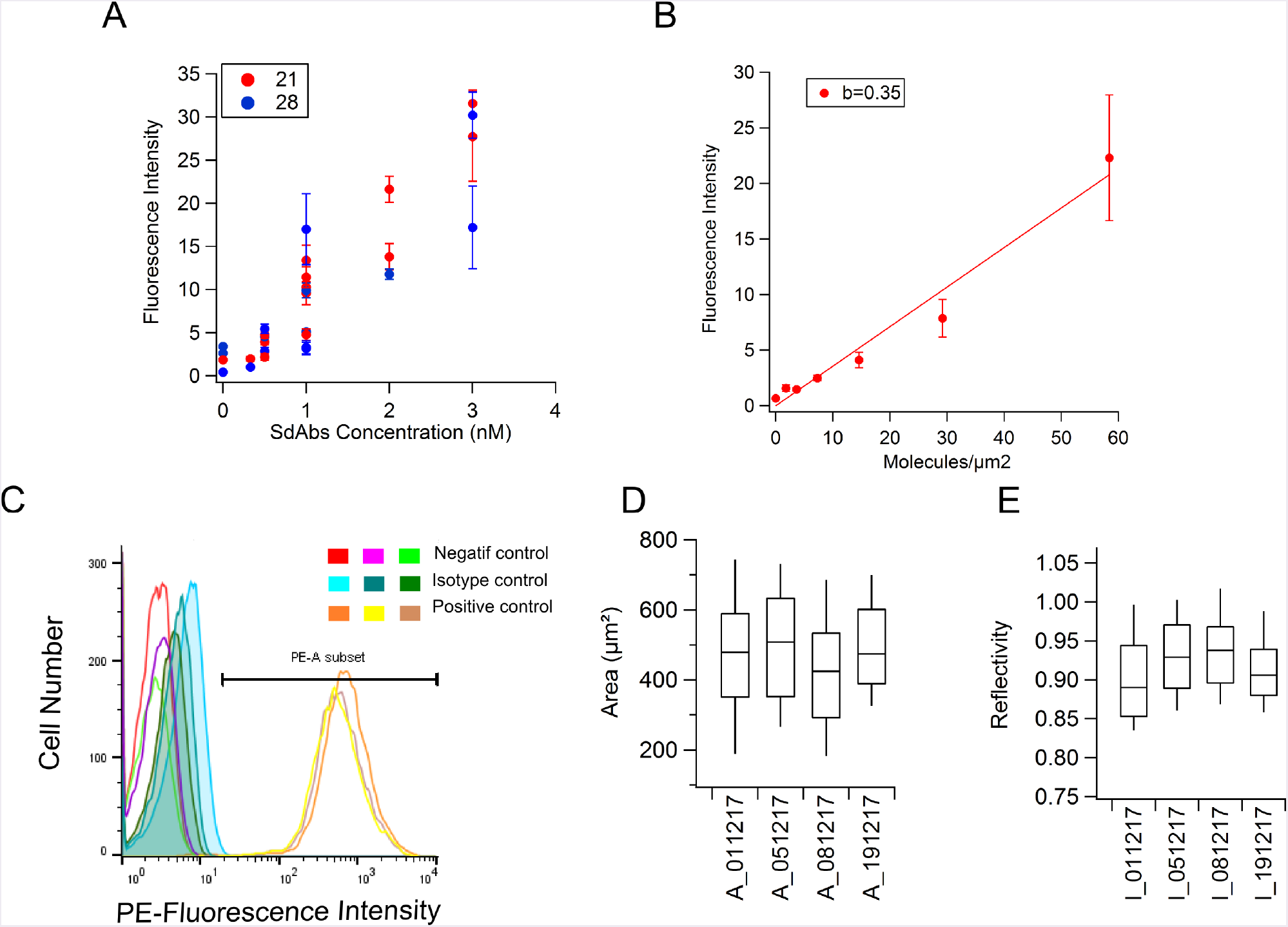
Controls of nanobody coated surfaces and NK92^hCD16^ cells. A) Fluorescence intensity values corresponding to the concentration of nanobody incubated on the Ibidi surface. (Fluorescence surface control was done at the end of the experiment). B) Calibration of the fluorescence intensity as a function of the surface density of nanobodies at the surface C) Fluorescence intensity histograms obtained by flow cytometry showing CD16 expression on NK cells. Superimposed positive curves indicate that CD16 expression is stable throughout all the period of cell culture. D,E) Distribution for four representative experiments of NK spreading area (D) and reflectivity (E) values obtained on surfaces coated with conventional anti-CD16 (clone 3G8), taken as a positive adhesion control. NK cell spreading with anti-CD16 coated surfaces was similar in all the experiments.

**Figure S5:**
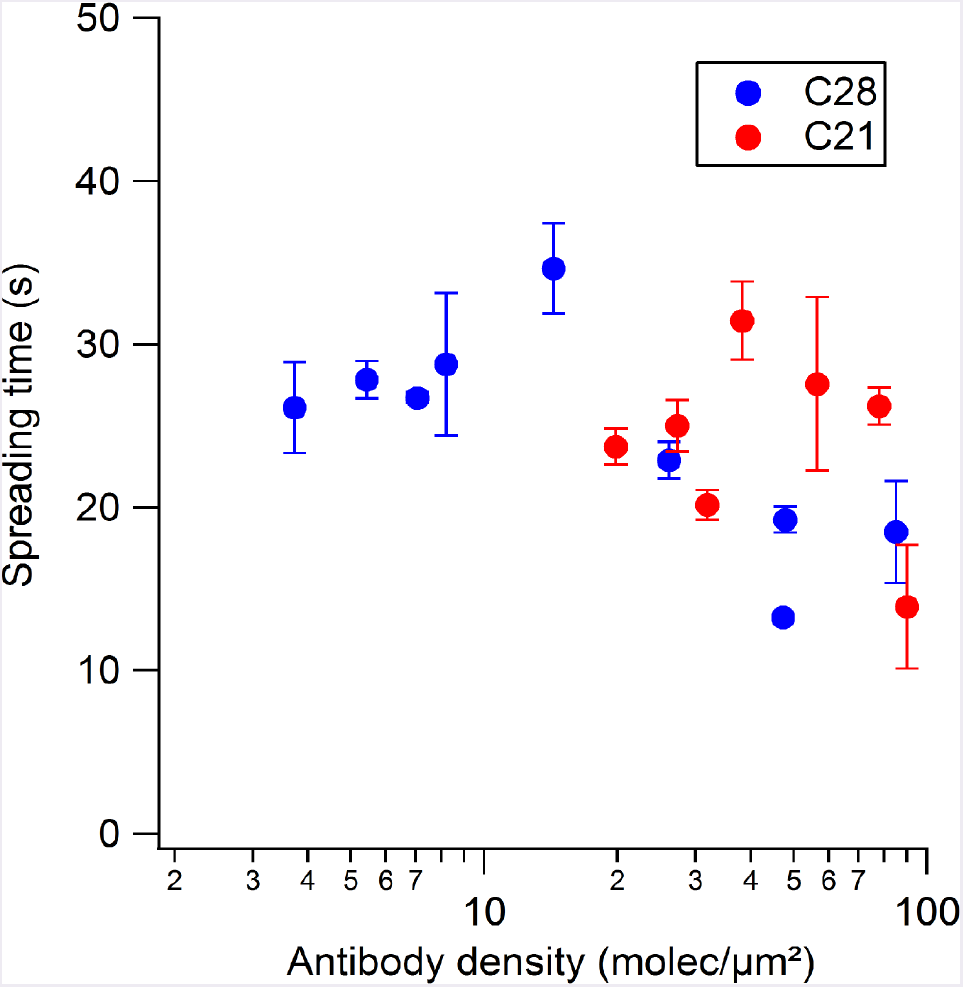
Cell Spreading kinetics. Individual NK cells engaging on surface coated with various densities of nanobodies were monitored over time with RICM. Spreading area versus time curves were fitted using a sigmoidal curve with time constant reported on the y axis. Each point and error bar represent the average and SEM of at least 10 cells.

**Figure S6:**
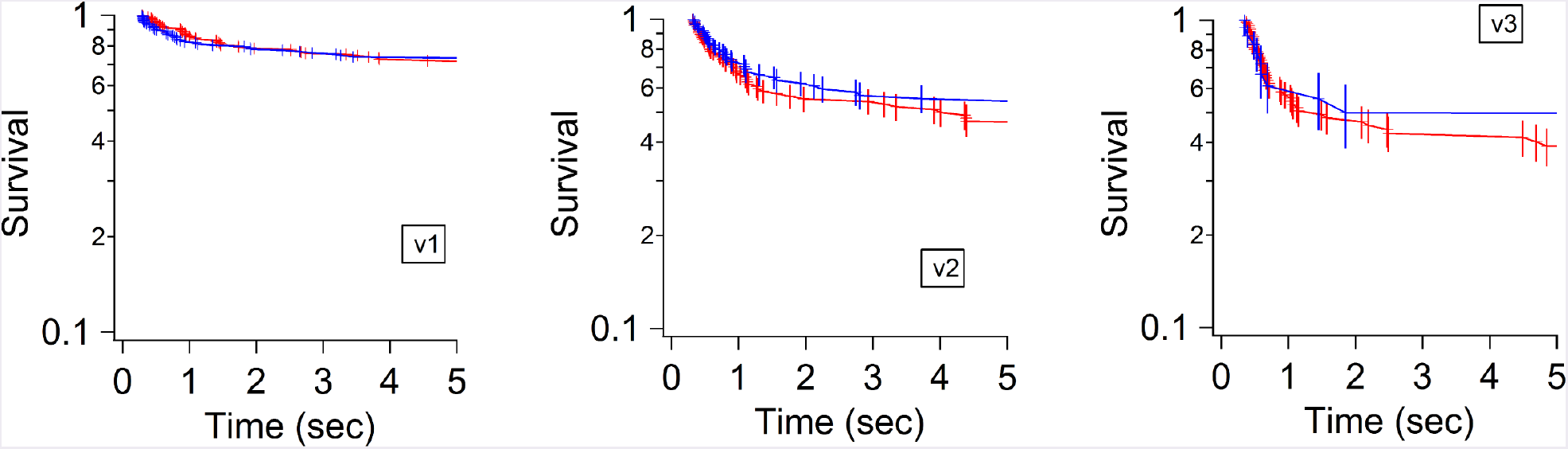
Survival curves of the transient adhesion of NK cells on nanobodies anti-CD16 coated surfaces (nanobodies density values on surface were between 6.5-12 molecules/*μ*m^2^) measured with the laminar flow chamber at shear stress of A) 0.075, B) 0.3 and C) 0.6 dyn/cm^2^. Red: C21; Blue: C28. Survival curves are built by the pool of arrested cells from at least three independent experiments.

**Figure S7:**
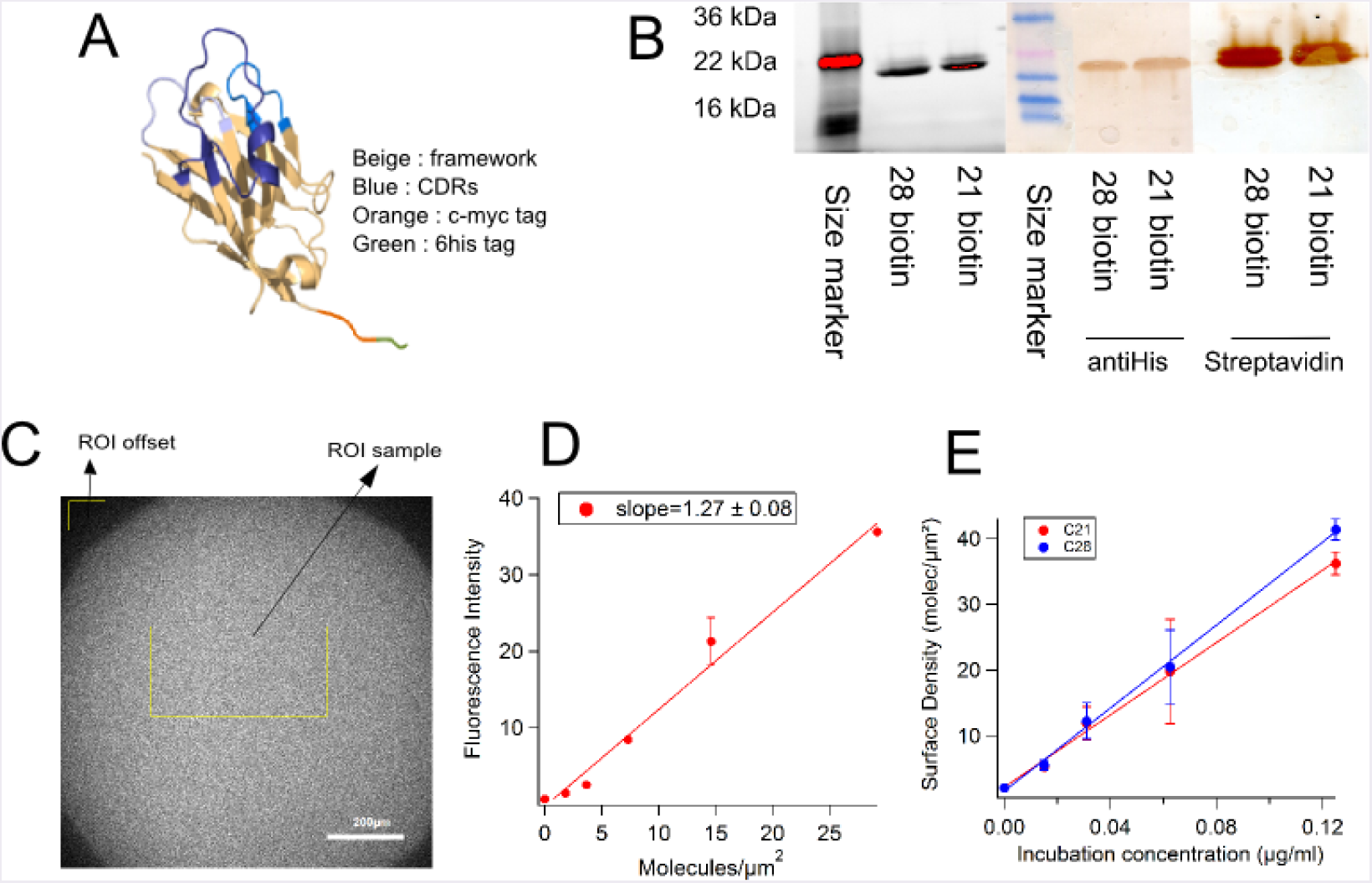
Nanobody structure and biotinylation process. A. Schematic representation of the nanobodies C21 or C28 with the corresponding His and c-Myc tag. B. Gel and Western Blot of C21 and C28 after biotinylation. On the gel, the strongest band corresponds to molecular weight of nanobodies (MW=15 kDa). On the Western Blot, anti His staining shows the presence of nanobodies C21 and C28 via the His tag and Streptavidin staining reveals the biotinylation of these nanobodies. C. Density of nanobodies on surfaces assessed using the detection of the Histag with a PE conjugated anti-His mAb. The image shows the fluorescence obtained after depositing 0.125 *μ*g/ml of nanobody C28 on the slide. The yellow rectangle visualizes the selected ROI for intensity measurement. D) Calibration curve giving the fluorescence intensity of anti-His-PE fluorescence antibody as function of the number of molecules/*μ*m^2^. The slope of the linear fit b=1.27 was used to determine nanobodies density. E) Graph showing the molecular density as function of the concentration of incubation of nanobodies. The density factor is the slope of the concentration *vs* density line.

**Figure S8:**
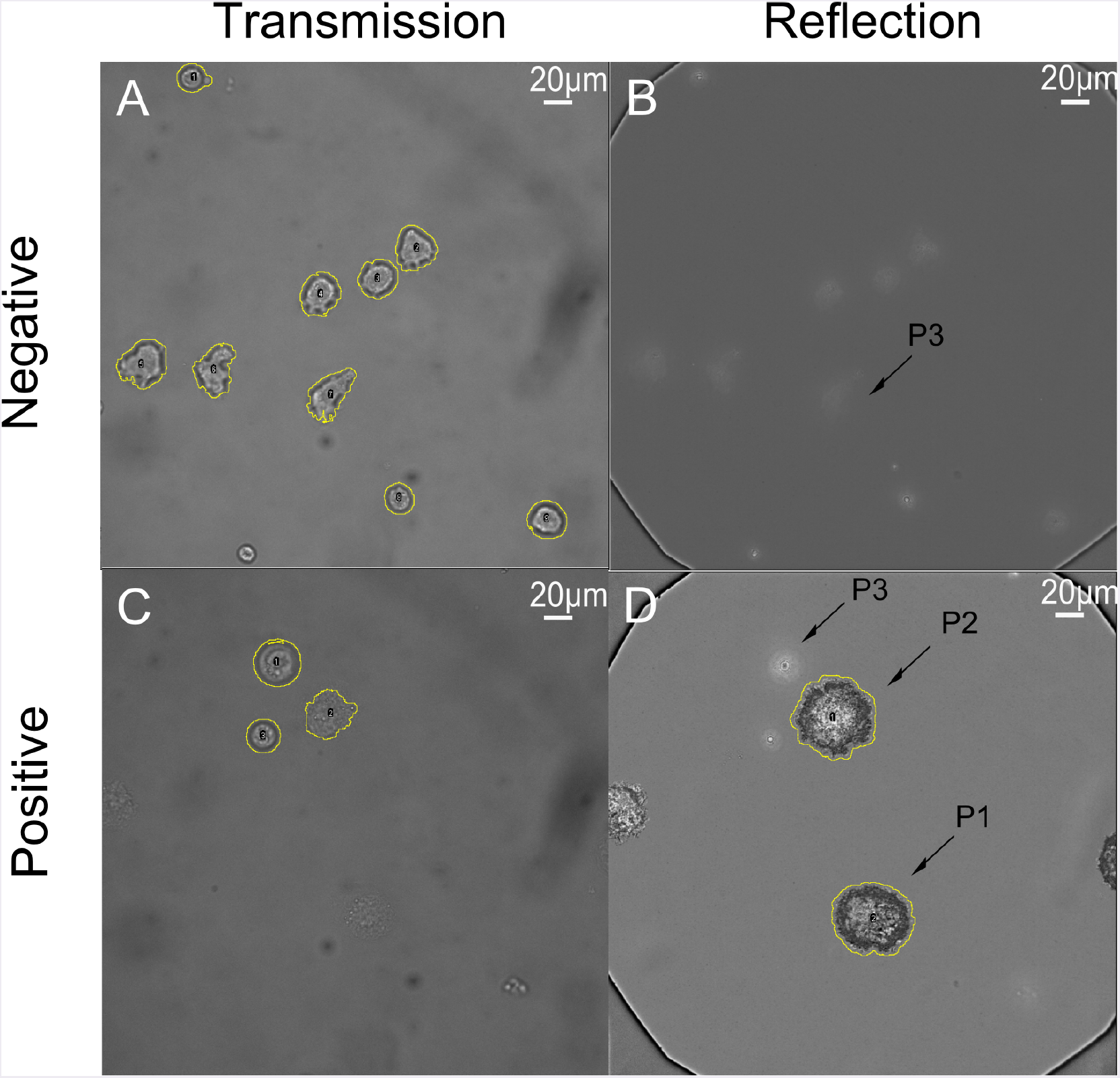
Images and procedure of segmentation of NK cells on anti-CD16 surfaces. A-B) Negative control (without nanobody on surface) showing the population P3 (no spread cells) as cells detected in transmission and not in reflection. C-D) Positive control, showing spread cells distributed among different subpopulations: P1 (detected only in RICM) and P2 (detected on both images) as well as non spread cells (P3).

**Figure S9:**
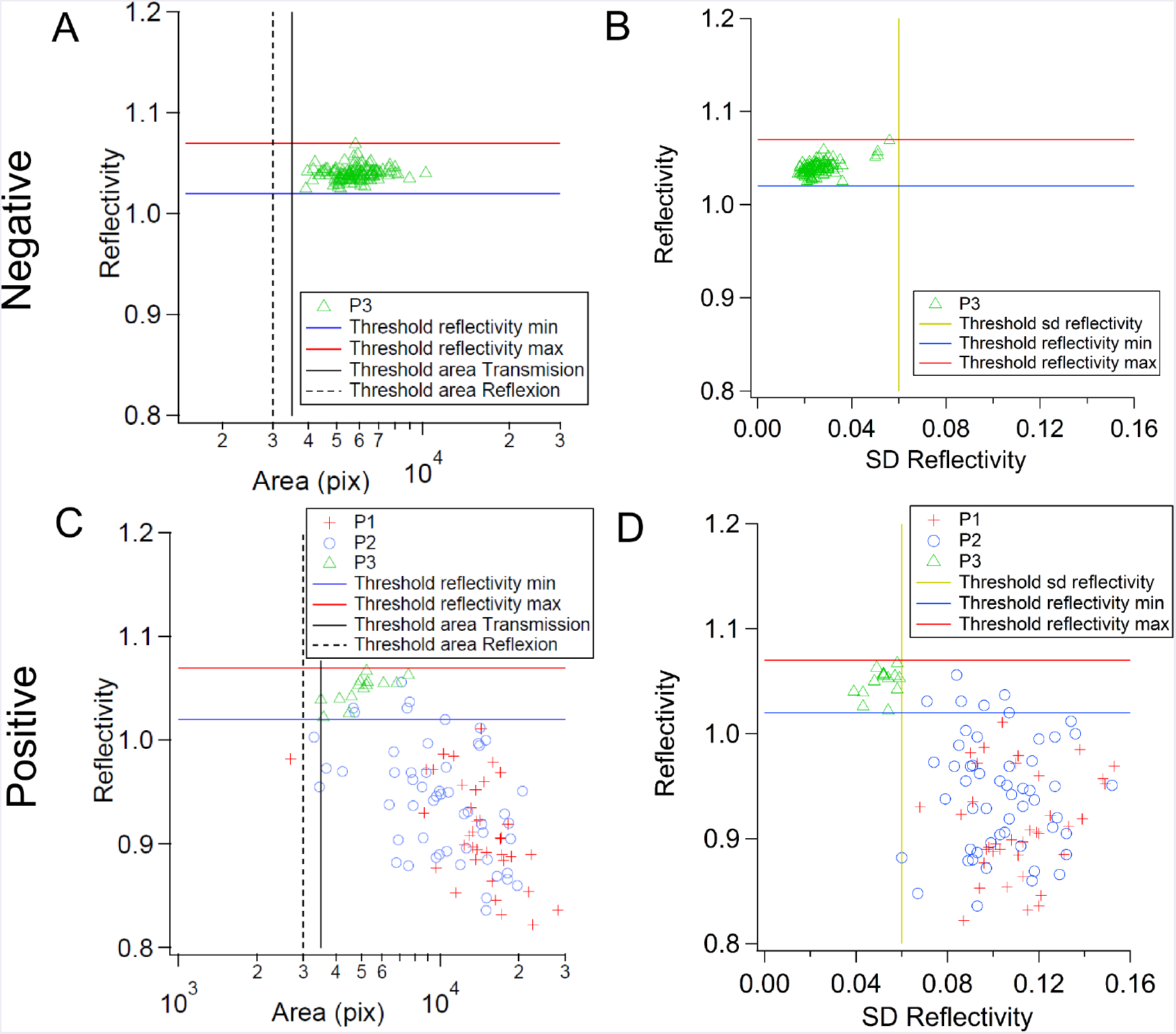
Graphs used to determine the parameters which defined the different populations. A. Graph of a negative control showing thresholds of area and mean reflectivity used for no adherent cells selection and P3 population between reflectivity thresholds. B. Graph of the same negative control showing the threshold of sd reflectivity and P3 population under the sd reflectivity threshold. C-D. Graphs of a positive control showing the thresholds of area, mean (C) and sd reflectivity (D) and the different adherent populations (P1 and P2) separated from the non adherent population (P3).

